# Phylotranscriptomics Reveals the Reticulate Evolutionary History of a Widespread Diatom Species Complex

**DOI:** 10.1101/2022.04.11.487918

**Authors:** Ozan Çiftçi, Andrew J. Alverson, Peter van Bodegom, Wade R. Roberts, Adrienne Mertens, Bart Van de Vijver, Rosa Trobajo, David G. Mann, Walter Pirovano, Iris van Eijk, Barbara Gravendeel

## Abstract

In contrast to surveys based on a few genes that often provide limited taxonomic resolution, transcriptomes provide a wealth of genomic loci that can resolve relationships among taxonomically challenging lineages. Diatoms are a diverse group of aquatic microalgae that includes important bioindicator species and many such lineages. One example is *Nitzschia palea*, a widespread species complex with several morphologically based taxonomic varieties, some of which are critical pollution indicators. Morphological differences among the varieties are subtle and phylogenetic studies on a few genes fail to resolve their evolutionary relationships. We conducted morphometric and transcriptome analyses of 10 *Nitzschia palea* strains to resolve the relationships among strains and taxonomic varieties. *Nitzschia palea* was resolved into three clades, one of which corresponds to a group of strains with narrow linear-lanceolate valves. The other morphological group recovered in the shape outline analysis was not monophyletic and consisted of two clades. Subsequent gene-tree concordance analyses and phylogenetic network estimations revealed patterns of incomplete lineage sorting and gene flow between intraspecific lineages. We detected reticulated evolutionary patterns among lineages with different morphologies and a resulting putative recent hybrid. Our study shows that phylogenomic analyses of many unlinked nuclear loci, complemented with morphometrics, can resolve complex evolutionary histories of recently diverged species complexes.

## Introduction

Diatoms (Bacillariophyta) are one of the most diverse and abundant groups of microalgae. Their ecological importance and high specificity towards different environmental parameters (e.g., salinity, pH, and nutrients) make diatoms ideal bioindicators (Smol and Stoermer 2010). Various environmental applications, such as water quality biomonitoring and palaeoecological reconstructions, rely on assessing the entire diatom community (Fritz et al. 1991, CEN 2004). Nearly all these applications are based on species identification using the famously character-rich and intricately ornamented siliceous cell walls of diatoms, which provide a wealth of characters for taxonomy and species delimitation. Quantitative morphometric methods have also been developed and proved reliable for distinguishing differences among diatom species (Theriot and Ladewski 1986, Pappas et al. 2014).

DNA sequencing has shown, however, that estimates of diatom diversity based on cell morphology are highly underestimated (Sarno et al. 2005, Sunagawa et al. 2015). An increasing number of molecular phylogenetic studies indicate that many diatom morphospecies comprises several distinct and often reproductively isolated lineages indistinguishable using morphological criteria alone, such as *Cyclotella meneghiniana* Kützing (Beszteri et al. 2005), *Pseudo-nitzschia delicatissima* Cleve (Hasle) (Quijano-Scheggia et al. 2009), *Gomphonema parvulum* Kützing (Kützing) (Kermarrec et al. 2013), and *Pinnularia borealis* Ehrenberg (Pinseel et al. 2019). Moreover, it is estimated that detectable genetic differences can arise between large populations of planktonic diatom species in just 10^2^ years (Lewis et al. 1997, Krasovec et al. 2019), and speciation can take place over periods of 10^3^ – 10^4^ years or less (Theriot 1992, Mann 1999).

Resolving relationships among recently diverged lineages is essential for improving our understanding of diatom systematics and biodiversity, recovering valuable information about their ecology and evolution, and ultimately improving environmental applications based on critical bioindicator species. In such cases, however, classical morphological characters traditionally used to separate species can be scarce or uninformative, while the unique features of taxonomically challenging taxa might be physiological or biochemical (Mann et al. 2021). Therefore, molecular phylogenies based on a broader genome sampling may provide better-resolved evolutionary relationships. Moreover, the expectation for recently diverged species is that paraphyletic or polyphyletic gene trees will reflect the speciation history, and the use of multiple unlinked loci from the nuclear genome is necessary to account for events such as incomplete lineage sorting (ILS) and gene flow (Mallet 2005, Alverson 2008). Transcriptomes are a valuable resource in this sense because they are relatively inexpensive to obtain, and their collection does not require a priori information about the target genome. Furthermore, tree-based orthology inference methods have been improved for nonmodel organisms and can now accommodate the complex nature of transcriptome data (Yang and Smith 2014, Emms and Kelly 2015, Cheon et al. 2020).

We use this approach to investigate *Nitzschia palea* (Kützing) W. Smith, a common bioindicator species with global distribution (Finlay et al. 2002). The nominate variety is described as tolerant to heavy metals and an indicator of pollution, while *Nitzschia palea* var. *debilis* (Kützing) Grunow prefers cleaner waters (Lange-Bertalot 1980, Van Dam et al. 1994, Sabater 2000, Potapova and Hamilton 2007). However, the type specimens of these varieties show considerable overlap in their morphological characters (Trobajo and Cox 2006). Morphometric, reproductive, and phylogenetic analyses demonstrated that it was impossible to separate the complex into the traditionally recognized varieties, although *N. palea* is a monophyletic group with several distinct lineages (Trobajo et al. 2009, Trobajo et al. 2010). Rimet et al. (2014) expanded these studies by including more global isolates and detected biogeographic signals but failed to find an objective criterion for marking varietal boundaries.

Here, we used *N. palea* as a model to infer the evolutionary history of a large diatom species complex for the first time by analyzing morphometric and transcriptomic data from 10 *N. palea* strains to (i) resolve the phylogenetic relationships between different lineages in the complex, (ii) investigate morphological characters congruent with molecular-based clades, and (iii) discriminate between ILS and gene flow to resolve patterns of reticulate evolution.

## Materials & Methods

### Sample collection and algal culturing

We acquired 10 *N. palea* strains from culture collections (Table 1). Four strains (DCG0091, DCG0092, DCG0094, and DCG0751) were cryopreserved in the collection. Three of these strains (DCG0091, DCG0092, and DCG0094) were among those studied by Trobajo et al. (2009, there referred to as Belgium3, Belgium1, and Belgium2) and TCC13901, TCC13903, TCC523, and TCC641 were included in the study of Rimet et al. (2014). All strains were grown in WC medium (Guillard and Lorenzen 1972) at 19 °C and on a 16 h light/8 h dark cycle. We routinely examined the live cultures under a Zeiss Axio Imager M2 microscope (Carl Zeiss BV, Breda, The Netherlands) and transferred the cells every 1–2 weeks based on the observed growth rates of individual strains.

**Table 1.**
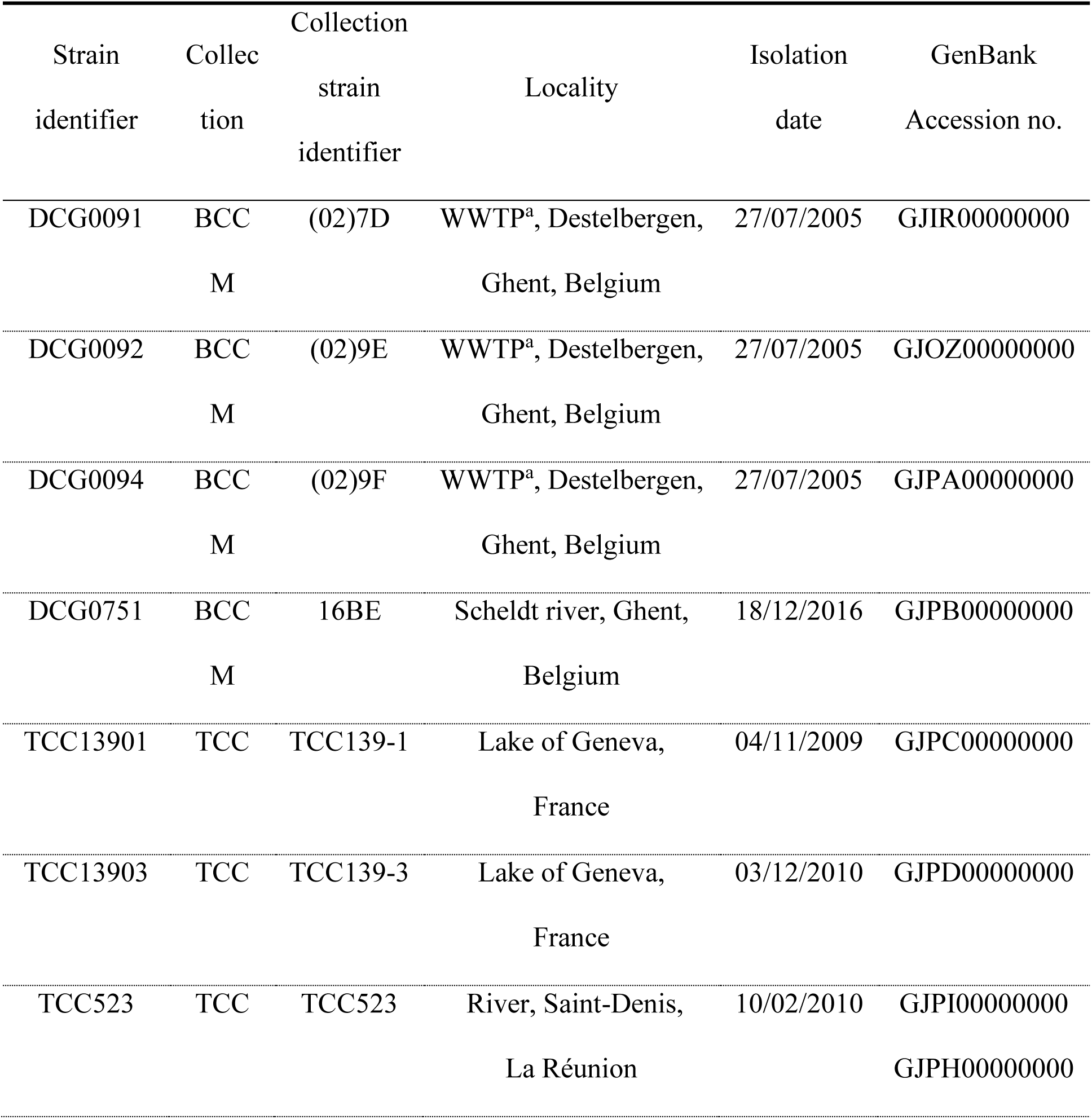
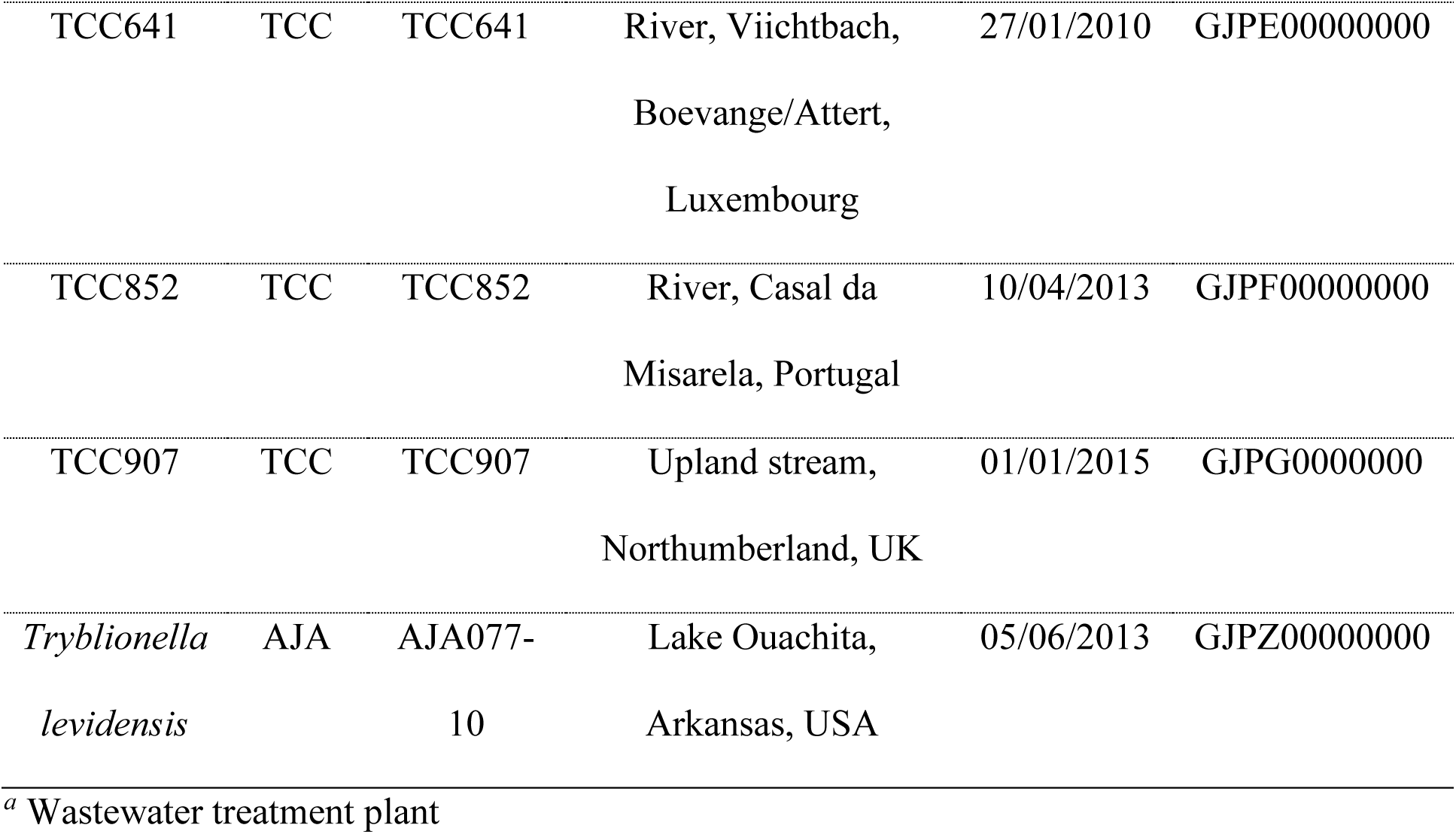
Collection, isolation information, and GenBank accessions of *Nitzschia palea* and *Tryblionella levidensis* strains analyzed (BCCM: Belgian Coordinated Collections of Microorganisms, Belgium; TCC: Thonon Culture Collection, France; AJA: Alverson Lab collection)

### SEM imaging, morphometrics, and shape outline analysis

Diatom cells were subcultured up to four times until harvesting for Scanning Electron Microscope (SEM) imaging. We oxidized the frustules with hydrogen peroxide and rinsed them several times with distilled water. Cleaned frustules were dried onto aluminum stubs and coated with a layer of platinum-palladium (Pt 80%, Pd 20%) in a Quorum Q150TS sputter coater (Quorum Technologies Ltd., UK). Four strains (DCG0091, DCG0092, DCG0094, DCG0751) were observed on a JSM-6480 Low Vacuum (JEOL) SEM platform at 10 kV, and the remaining strains were observed on a JSM-7600F Field Emission (JEOL) Scanning Microscope system at 5 kV. We collected 50 images per strain at magnifications ranging from 8000–15000x. The width (µm), length (µm), stria density (per 10 µm), and fibula density (per 10 µm) of each image were measured or counted using the software package Fiji-ImageJ (Schindelin et al. 2012). Measurements of stria and fibula densities were made in the center of the valve along its apical axis over 10 µm. Additional descriptive morphological data investigated in previous studies on *N. palea* were also recorded (Trobajo & Cox, 2006). For analysis of shape outlines, we used Elliptic Fourier Analysis (EFA) which is also implemented in diatom morphometric tools such as DiaOutline (Wishkerman and Hamilton 2018) and SHERPA (Kloster et al. 2014). We extracted valve outlines from SEM images using the “Quick Selection” tool of Adobe Photoshop CC 2019 and exported these on a white background. We used the R package Momocs (Bonhomme et al. 2014) for (i) extracting x and y coordinates of the valve outline shape, (ii) pre-processing (i.e., smooth, center, scale, and align), and (iii) computing Elliptic Fourier Transforms (EFT). Finally, we performed a Principal Component Analysis (PCA) analysis using Momocs and visualized the first two principal components (PC). Boxplot and PCA figures were produced using the R package ggplot2 (Wickham and Chang 2016).

### RNA extraction and transcriptome sequencing

We extracted total RNA from exponentially growing cultures using the Qiagen RNeasy Plant Mini Kit (Qiagen Benelux BV, Venlo, The Netherlands). First, we removed the excess culture medium and concentrated the cells through repeated centrifugation steps at 2900x*g* for 10 min.. Final suspensions were transferred to 2 ml tubes containing 0.5 mm zirconia/silica beads (BioSpec, Lab Services BV, Breda, The Netherlands) and the lysis solution provided with the kit. Cells were lysed using a Qiagen Tissue Lyser II (Qiagen Benelux BV, Venlo, The Netherlands) by bead beating for 3 minutes. Subsequent steps followed the manufacturer’s protocol. We assessed the quantity and quality of RNA samples on an Agilent 2100 Bioanalyzer system (Agilent Technologies, Amstelveen, The Netherlands), and RIN values ranged from 5.2 to 7.2. Sequencing libraries were prepared using the Illumina TruSeq Stranded mRNA Library Preparation Kit (Illumina, The Netherlands). Sequencing was performed at Baseclear BV (Leiden, The Netherlands) in two separate runs on a NovaSeq 6000 platform (Illumina), generating 2×150 bp paired-end reads. RNA extraction and transcriptome sequencing of the outgroup species, *Tryblionella levidensis* W. Smith, followed Parks et al. (2018). Raw sequencing reads were deposited in the Sequence Read Archive (SRA) database of the National Center for Biotechnology Information (NCBI) under BioProject accession PRJNA756685.

### Transcriptome assembly

Filtering and assembly of sequencing reads followed Parks et al. (2018) with minor modifications. We inspected the quality of the raw reads using SolexaQA++ (ver. 3.1.7) (Cox et al. 2010). Error-correction was performed using BFC (Li 2015) with parameters: -k 31 (k-mer length), -s 50. Sequencing reads were trimmed using Trimmomatic (ver. 0.39) (Bolger et al. 2014) with the options ‘ILLUMINACLIP:2:30:10 LEADING:3 TRAILING:3 SLIDINGWINDOW:4:12 MINLEN:50’. As the final step of the pre-assembly process, we removed all reads mapping to plastid and mitochondrial genomes of *N. palea* (strain NIES-2729; Kamikawa et al. 2018, AP018511 and AP018512, respectively) and the SILVA rRNA database using bowtie2 (ver. 2.4.1) in “--very-sensitive-local” mode (Langmead and Salzberg 2012). The filtered reads were assembled using Trinity (ver. 2.10) with strand-specific options (Grabherr et al. 2011). We assessed assembly quality by checking the recovery of conserved eukaryotic orthologs in the BUSCO database (ver. 4) (Waterhouse et al. 2018). Assembled nuclear transcripts were translated into amino acid sequences with TransDecoder (ver. 5.5.0) (Haas 2020) with guidance from BLASTP (ver. 2.8.1) searches against the SwissProt database (The Uniprot Consortium 2019) with an e-value of 1e^-3^, and HMMER (Finn et al. 2011) searches against the Pfam database (El-Gebali et al. 2019). Redundant transcripts were filtered using CD-HIT (ver. 4.8.1) (Li and Godzik 2006) with a sequence identity threshold of 0.99 and a word length of five. Finally, we quantified transcript abundances using Salmon (ver. 1.0.0) (Patro et al. 2017) and selected the most highly expressed isoform per trinity gene for downstream analyses.

### Orthology inference and species tree reconstruction

We used OrthoFinder (ver. 2.4.0) (Emms and Kelly 2019) to infer putative orthologous clusters from the complete set of predicted proteins. Building and pruning homolog trees for species tree reconstruction followed the *phylogenomic dataset construction* pipeline of Yang and Smith (2014). We extracted coding sequences for orthogroups that contained at least one transcript per strain, aligned the amino acid sequences with MAFFT (ver. 7.47) (Katoh and Standley 2013), and trimmed alignments with Phyx (Brown et al. 2017) using a minimal column occupancy threshold of 0.2. Orthogroup gene trees were constructed using RAxML (ver. 8.2.12) (Stamatakis 2014) with the GTRCAT model and 100 rapid bootstrap replicates. Outlier long branches were removed with TreeShrink (Mai and Mirarab 2018), and mono- and paraphyletic tips that belonged to the same taxon were pruned, along with deep paralogs, using respective Python scripts from Yang and Smith (2014). Tips with the largest number of unambiguous characters in the trimmed alignment were retained, and these were realigned with MAFFT. These amino acid alignments were used to guide codon alignments with pal2nal (Suyama et al. 2006). These alignments were again trimmed using Phyx as described above, and maximum likelihood homolog trees were inferred using IQ-TREE 2 (Minh et al. 2020a) with 100 bootstrap replicates and the TESTMERGE procedure with codon position partitions specified for each alignment. TESTMERGE selects the best-fit partitioning scheme followed by model selection and tree construction. As the last step of the pruning pipeline, we extracted one-to-one orthologs from these bootstrapped homolog trees using Yang and Smith’s (2014) MI strategy with a long internal branch cutoff of 0.6. Ortholog trees were inferred using the same strategy with homolog tree inference. Finally, we estimated two species trees using summary-coalescent and concatenation-based approaches. We used ASTRAL-III (Zhang et al. 2017) with the set of unrooted ortholog trees to estimate the first species tree with the support values calculated as the local posterior probability (LPP). The concatenated ML analysis was performed using IQ-TREE 2 with 100 bootstrap replicates, where support values are calculated as the bootstrap support (BS). The model of sequence evolution for each locus was calculated using ModelFinder, as implemented in IQ-TREE 2, with separate substitution models and separate evolutionary rates across sites. A summary workflow for all steps from data collection to species tree estimation is given in Figure S1. *Tryblionella levidensis* was specified as the outgroup taxon in the ASTRAL-III analysis, and the ML species tree was rooted manually using FigTree (ver. 1.4.4).

### Quantifying genealogical concordance

Gene concordance factors were computed using IQ-TREE 2 by comparing the alternative resolutions of quartets of taxa around each branch (Minh et al. 2020b). The gene concordance factor (gCF) describes the proportion of concordant gene trees with a given split in the species tree, whereas the proportions of concordant gene trees supporting the two alternative topologies are reported as gene discordance factors, gDF_1_ and gDF_2_. The proportion of all other conflicting resolutions, which result in different arrangements of paraphyletic gene trees, is reported as the third gene discordance factor, gDF_P_. If ILS is the source of the discordance, gene trees supporting the two alternative topologies (gDF_1_ and gDF_2_) are expected to occur in roughly equal frequency (Huson et al. 2005) and the significance of this prediction was tested using a chi-squared test. Alternatively, unequal proportions of gene trees might indicate gene flow between clades (Huson et al. 2005). Under strict ILS assumptions (and ignoring gene tree error), we expect the two discordant topologies to occur in equal frequency (Huson et al. 2005). Gene concordance and discordance factors were mapped onto the species tree as pie charts, and the mean shapes for each strain were illustrated at the tips using the R packages ape (Paradis et al. 2004) and ggtree (Yu et al. 2017).

### Phylogenetic network estimations and tests for introgression

We estimated phylogenetic networks while accounting for ILS and gene flow using two approaches to explore taxon relationships. First, we estimated a phylogenetic network under maximum pseudo-likelihood using the InferNetwork_MPL command in PhyloNet (Yu and Nakhleh 2015, Wen et al. 2018). We used the complete set of rooted gene trees, applied a bootstrap support threshold of 95, ran ten independent network searches, and sequentially tested the allowance of 0 to 3 hybridization/reticulation branches. We selected the run with the highest log pseudo-likelihood as the best estimate. A sharp improvement in score is expected until it reaches the best value and has a slower, linear improvement after that. We visualized PhyloNet results using Dendroscope (Huson et al. 2007). Second, we estimated a phylogenetic network using the NANUQ algorithm (Allman et al. 2019). We used all gene trees and ran NANUQ through the MSCquartets R package (Rhodes et al. 2021). We used a small alpha (0.01) and a large beta (0.95) for hypothesis testing following the developer’s recommendations. We used the test results on the quartet counts from all gene trees to calculate a network distance matrix between taxa. We then inferred a split network from the distance matrix using the Neighbor-Net algorithm (Huson and Bryant 2006). As a complement to the network approaches, we also performed introgression tests for the putative hybrid (TCC907) with Patterson’s *D*-statistic (Durand et al. 2011), commonly known as the ABBA-BABA test. The ABBA-BABA test is based on site patterns in the alignments under a null ILS model (i.e., no gene flow). Given the genome sequences of three ingroups and an outgroup population with the relationship (((P1,P2),P3),O), ABBA sites are those at which P2 and P3 share a derived allele (B), while P1 has the ancestral state (A). The BABA pattern represents sites at which P1 and P3 share the derived state. The ABBA-BABA test approximates the proportion of the genome represented by these two discordant topologies. In the absence of any deviation from a strict bifurcating topology (i.e., no gene flow), we expect to find roughly equal proportions of ABBA and BABA site patterns in the genome. The *D*-statistic test is used to quantify deviations from this proportion. We calculated *D* using the calcD function in the R package evobiR (Blackmon and Adams 2015) and assessed the significance of *D* using a block jackknife approach (block-size 1000 sites, 1000 replicates).

## Results

### Morphometric data and valve shape outlines

The valves that we examined with SEM had the following morphological features: (i) linear-lanceolate to lanceolate valve outlines with rostrate to subcapitate apices, (ii) irregularly spaced fibulae with median fibulae not more widely spaced, (iii) no central raphe endings, (iv) terminal raphe fissures that internally end in helictoglossa (Figure 1, E1), and (v) polar raphe endings that turn towards the same side in a single valve (Figure 1, F1 and F2). These characters agree well with the analysis of the type material of *N. palea* from the Kützing collection at the Natural History Museum, London, UK (Trobajo & Cox, 2006). The only exceptions were valves with deformed features.

**Figure 1.**
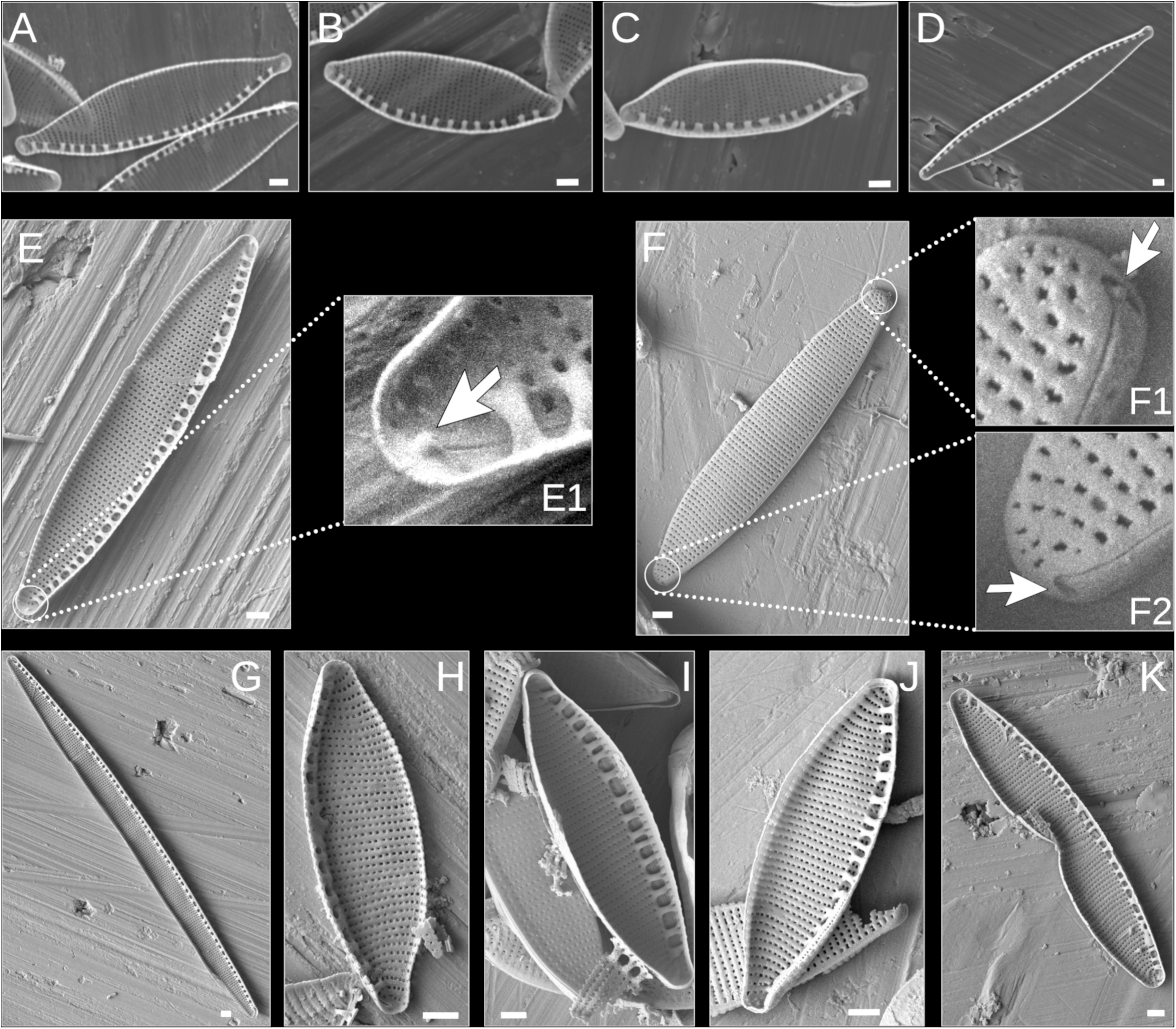
Scanning electron micrographs of *Nitzschia palea* strains analysed in this study. (A) DCG0091, (B) DCG0092, (C) DCG0094, (D) DCG0751, (E) TCC13091, (E1) Lower apex of TCC13901 showing helictoglossa, (F) External view of TCC13901, (F1) Upper apex of TCC13901 showing the polar raphe ending, (F2) Lower apex of TCC13901 showing the polar raphe ending, (G) TCC13903, (H) TCC523, (I) TCC641, (J) TCC852, (K) TCC907, representing a teratological form. All scale bars equal 1 µm.

The observed morphometric measurements, summarized in Figure 2, overlap with the ranges indicated for the types of *N. palea* (Trobajo & Cox, 2006), except for deformed valves and the mean lengths of five strains that were small due to size reduction caused by long-term cultivation (Figure 1K; Table S2). Specifically, most of the cells in strain TCC907 had striae and fibulae deformations and retractions at the center of the valve margins on one side. We removed this strain from the shape outline analysis and only used its valve lengths and widths. Many valves of TCC641 and TCC523 had striae and fibulae deformations, and we measured a minimum of 30 valves for these two strains. The number of images used in the EFA for TCC641 and TCC852 was lower due to valve outline deformities (Table S1). The deformations of TCC907 and TCC641 are apparent in Figure 2, with low stria densities that fall out of the ranges described from the type specimens (Table S2, Trobajo & Cox, 2006). We observed two different length ranges (greater than 34.5 µm and less than 25 µm) for TCC13903 (Figure 2). This strain is known to self-reproduce based on long-term observations (F. Rimet, personal communication, March 29, 2021). Its stria density varied least among all strains analyzed, providing corroborative evidence for the presence of a single taxon with two different length ranges, possibly representing different stages of its life cycle (Table S2).

**Figure 2.**
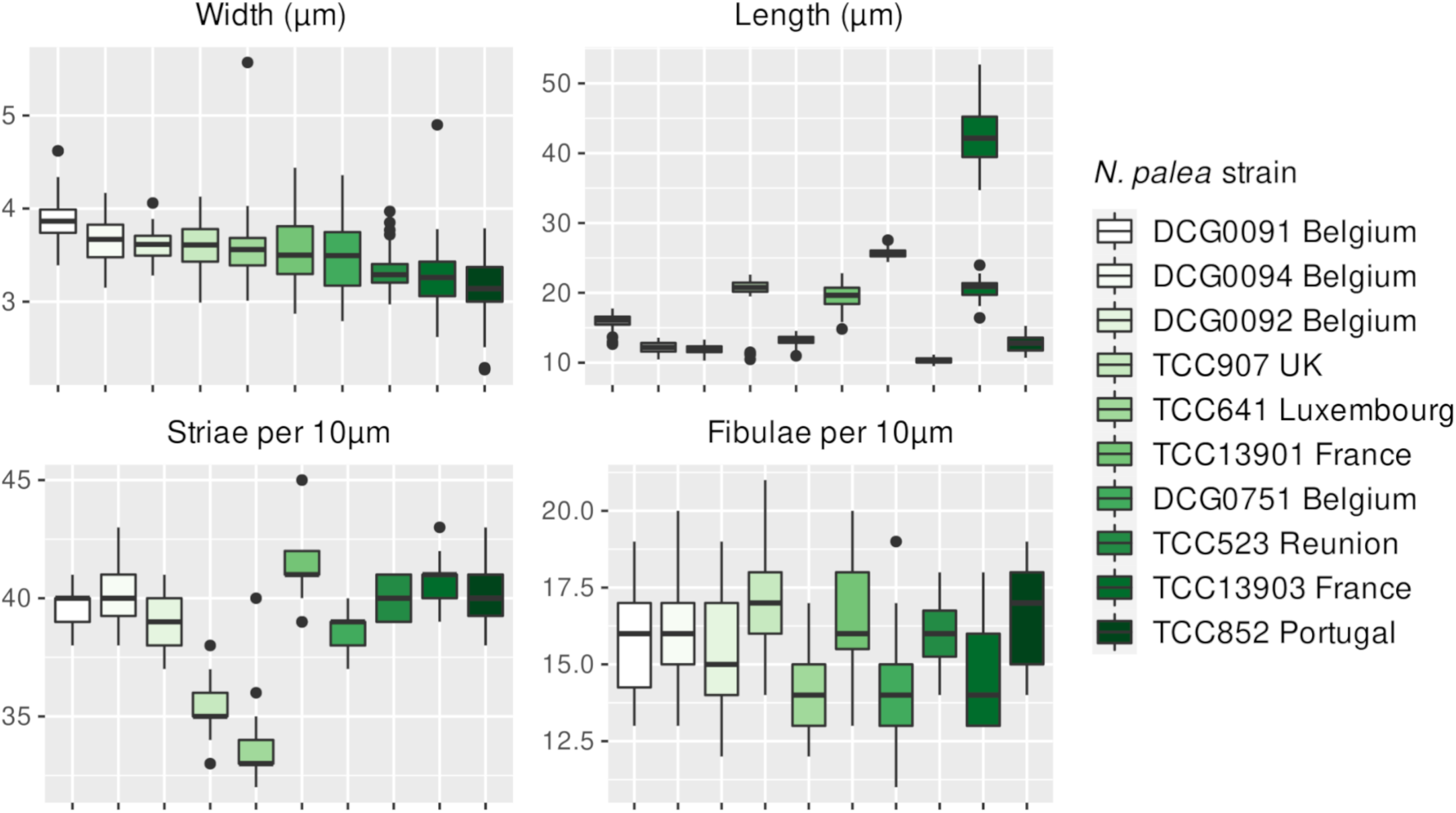
Boxplot summary of morphological measurements of *Nitzschia palea*. Black dots represent outliers. Strains are ordered based on their mean width.

Based on the morphological criteria used in Trobajo et al. (2009), none of the strains in this study can be assigned to *N. palea* var. *debilis* (i.e., less than 3.5 µm wide with stria density above 43 per 10 µm). Four strains (DCG0751, TCC523, TCC13903, and TCC852) were narrower than 3.5 µm. However, the highest mean stria density in this group was 40.68 per 10 µm (TCC13903) (Table S2). Two out of these four narrow strains (TCC523 and TCC852) had many deformities. The strain with the largest mean stria density in our sample set was TCC13901 (41.19 per 10 µm) with a mean width of 3.53 µm. The first principal component explain 98.2% of the total variation in valve shape (Figure 3A) and showed a gradient from lanceolate to linear-lanceolate outlines (Figure 3B). Two narrow strains without deformities (TCC13903 and DCG0751) and the strain with the largest mean stria density (TCC13901) overlapped at the linear-lanceolate end of the morphospace (Figure 3A). The remaining six strains included in the shape outline analysis (TCC523, TCC641, TCC907, DCG0091, DCG0092, and DCG0094) were much shorter than the ranges described for *N. palea*, probably due to size reduction caused by long-term cultivation. DCG0092 had the lowest mean stria and fibula densities in this group, whereas TCC852 had the largest (Figure 2). These six strains were positioned at the lanceolate end of morphospace in shape outline analysis, showing a slight overlap with the linear-lanceolate group (Figure 3A). The second principal component in the shape outline analysis explained less than 1% of the total variation in valve shape outline (Figure 3A) but showed a gradient from subcapitate to rounded ends (Figure 3B). All strains overlapped considerably based on this axis, and there was no apparent clustering.

**Figure 3.**
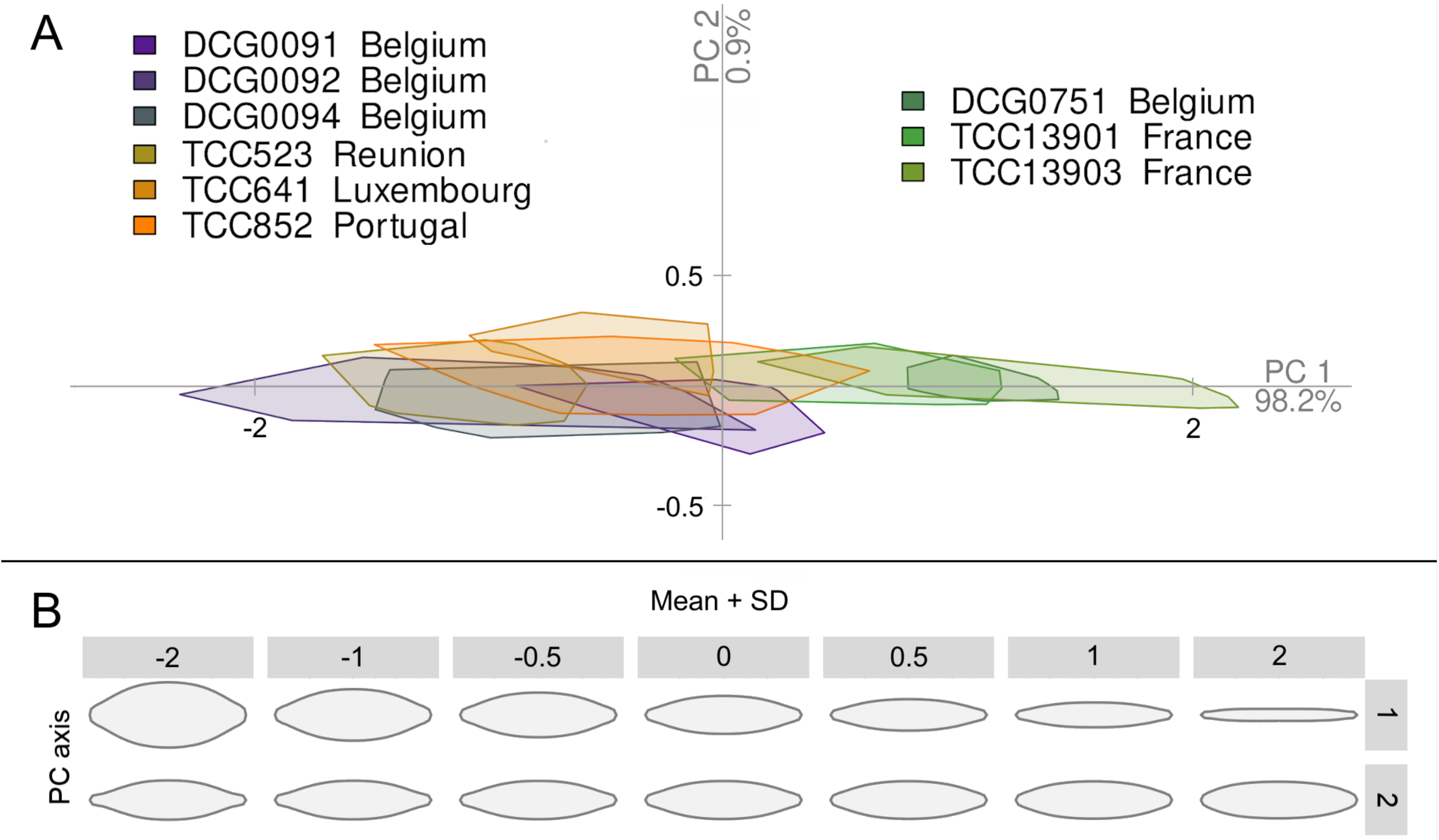
(A) PCA results of *Nitzschia palea* valve outline analysis. (B) Shape variation along the first two principal component axes. Standard deviations were calculated as the square roots of the eigenvalues of the covariance matrix

### Transcriptome assemblies, orthology inference, and species tree estimation

We assembled transcriptomes for 10 *N. palea* strains and an outgroup, *T. levidensis*. Datasets ranged in size from 15.6 to 33.4 million reads per strain. Two sets of reads were assembled for strain TCC523, with duplicates subsequently masked during the tree pruning procedure. Trinity assemblies ranged in size from 23791 to 110789 genes and 34388 to 157240 transcripts, including isoforms. The average BUSCO recovery of the assembled transcriptomes was 74 ±14%, indicating that we captured a large portion of the gene space in each strain. After filtering to remove transcripts with low expression levels, the final dataset used for ortholog clustering contained 16109–62583 trinity transcripts per strain. Among the 81016 total orthogroups identified by OrthoFinder, 1260 included data for all ten strains. We obtained 515 homolog alignments with complete-taxon-occupancy by estimating trees from these orthogroups, cutting long internal branches, and trimming and masking monophyletic tips (Yang and Smith 2014). Using Yang and Smith’s (2014) MI strategy, a final tree-pruning step to select orthologs resulted in 183 one-to-one ortholog trees with alignment lengths ranging from 396 to 4935 bp. The total concatenated alignment length for these 183 orthologs was 283107 bp, where 137025 (48.4 %) of these sites were constant, 65405 (23.1 %) were parsimony informative, and 80677 (28.5 %) were variable but parsimony uninformative. The topologies of ASTRAL and ML species trees were identical. Three main clades (Figure 4, clades A–C) were recovered with maximum LPP and BS values (LPP=1, BS=100) with both methods (Figure 4). All other branches in the estimated species trees, except for the internal branch in Clade C (LPP=0.8, BS=55), were strongly supported (LPP>0.96, BS=100).

**Figure 4.**
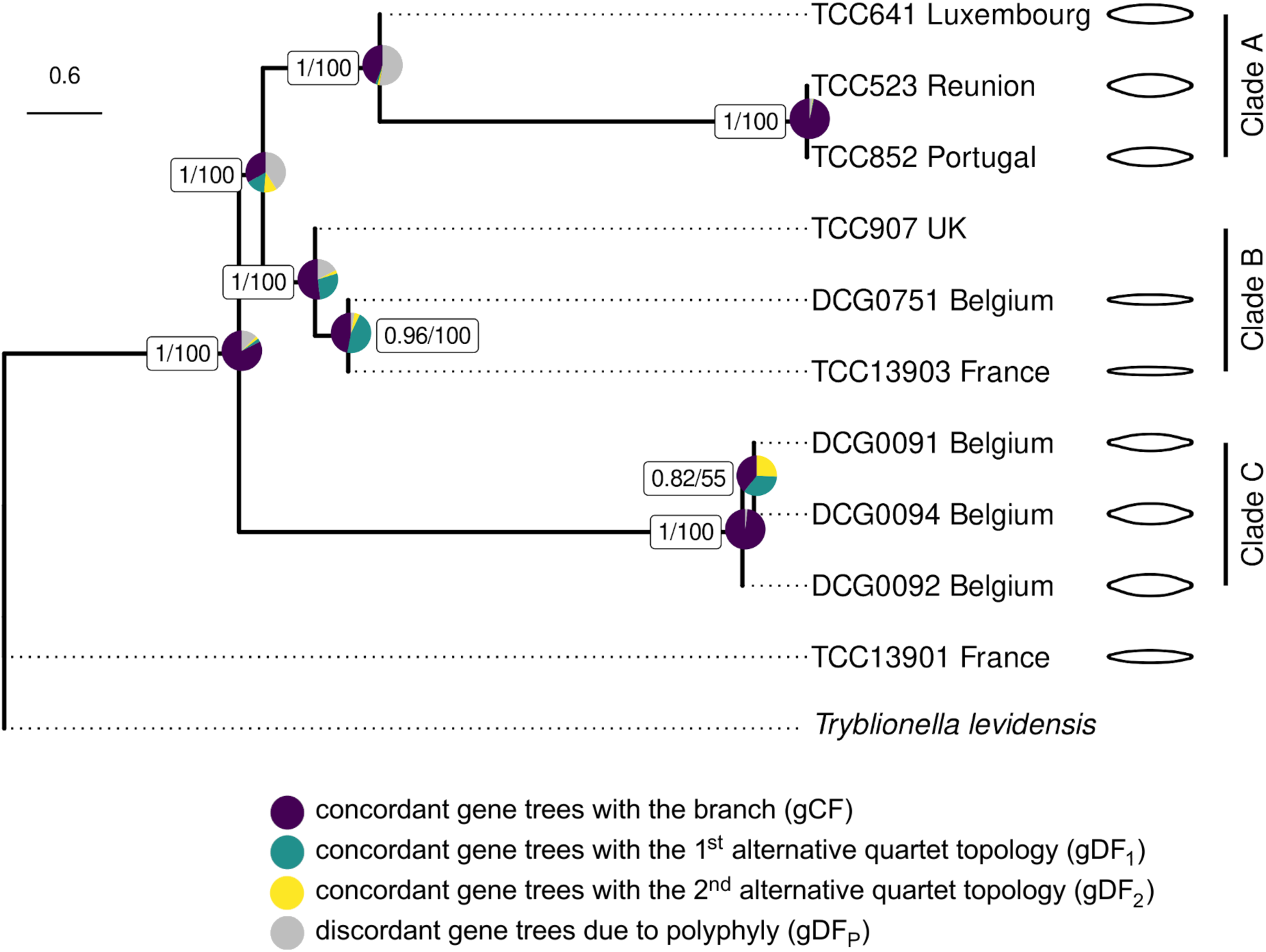
ASTRAL-III species tree estimated from 183 orthologs. Pie charts show IQ-TREE 2 gene concordance and discordance factors. ASTRAL-III LPP values (left) and bootstrap support from concatenated ML analysis (right) are given in boxes. Branch lengths are in coalescent units. Dotted lines are used to align the tip labels. Mean shapes from EFA for each strain are illustrated at the tips. No outline image is given for TCC907, which had deformed outlines representing an artificial shape due to culture conditions.

### Gene tree concordance analyses, phylogenetic network estimations, and tests for introgression

Among recently diverged taxa, individual gene trees can disagree with the underlying species tree due to ILS or gene flow (Maddison 1997, Degnan and Rosenberg 2006). We obtained maximum support values for the relationships among the three main clades in the species trees, despite evidence for widespread ILS in the evolutionary history of the *N. palea* species complex (Figure 4). In our gene tree concordance analysis, only 61 of the 183 gene trees supported a sister relationship of clades A and B (Figure 4). A large proportion of the discordance for this relationship was due to polyphyly (41%), indicating poor support in the gene trees for this split, and the proportions of gene trees with alternate topologies gDF_1_ (15%) and gDF_2_ (11%) were not significantly different for this branch (Figure 4), consistent with ILS. The split between TCC641 and the other two Clade A strains also was poorly supported (gDF_P_=52.46%). Deeper in the species tree, a large majority (151) of the gene trees supported a sister relationship of Clade C to the common ancestor of clades A and B (Figure 4). Chi-squared tests for unequal gDF_1_ and gDF_2_ values were significant (*P* < 0.05) only for the two branches in Clade B, suggestive of a non-ILS process occurring here.

We detected large proportions of gene trees concordant with the first alternative topologies for both nodes in Clade B (Figure 4, green vs. yellow), consistent with gene flow between Clade A and Clade B populations. These inferences were based on phylogenetic analyses that fit a model of strict bifurcation. To test for the possibility of a more complex network model, we used PhyloNet to estimate the best network from the set of 183 gene trees. This analysis identified TCC907 in Clade B as a putative hybrid between Clade A and DCG0751 (Figure 4, Clade B) with near-equal γ values (inheritance probabilities) (Figure 5A). To further test the hypothesis that TCC907 is of hybrid origin, we used NANUQ and MSCQuartets to estimate a phylogenetic network, and this analysis revealed further evidence for reticulation between Clade A and Clade B populations, with strain TCC907 placed as an intermediate between these parent clades (Figure 5B). The ABBA-BABA test for the putative hybrid, TCC907, was performed using the concatenated alignment of 183 orthologs where three Clade A strains were specified as one parental population and the other two members of Clade B as the second. Our results revealed a substantial excess of ABBA site patterns with a significant positive *D*-statistic regardless of the order of parental populations, further corroborating the results from the network analyses (D: 0.6/0.9; Z-score: 87/630).

**Figure 5.**
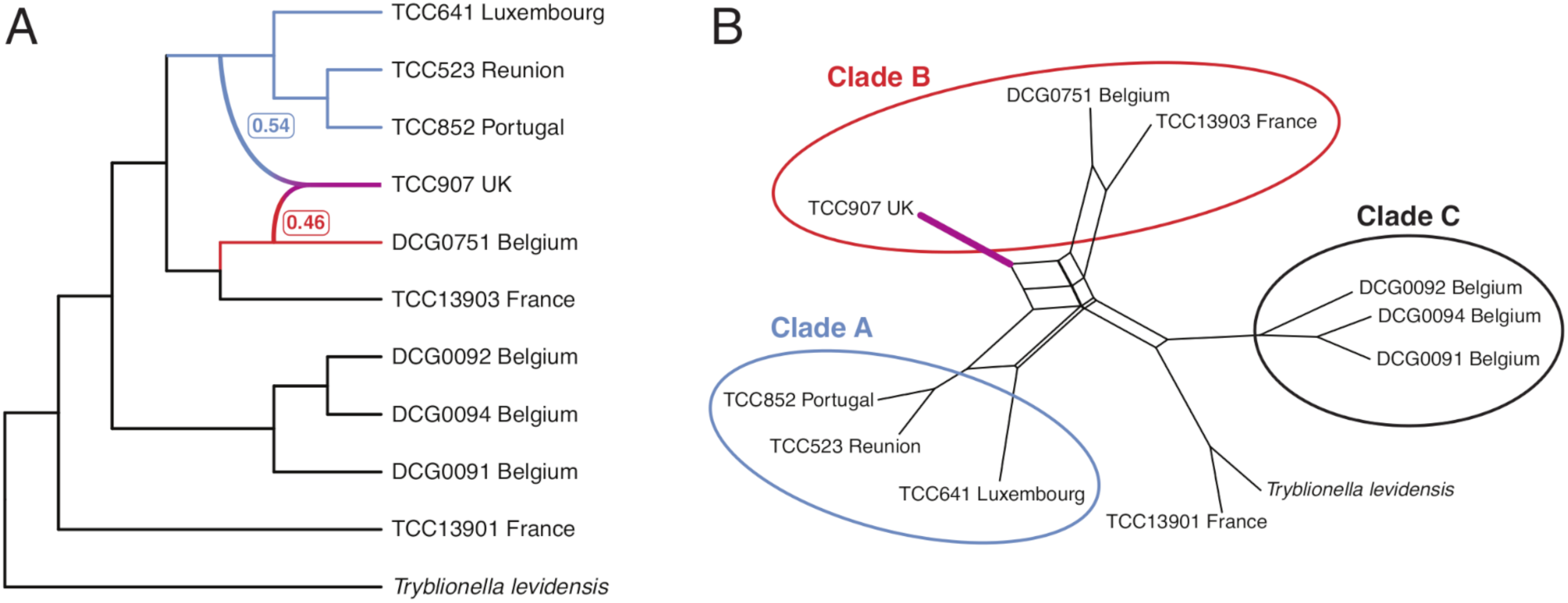
Phylogenetic networks estimated using (A) PhyloNet with inheritance probabilities (γ) from parent lineages of the putative hybrid indicated in *Nitzschia palea*, and (B) hybrid detection with NANUQ. Clade labels correspond to those recovered in the species tree (Fig. 4).

## Discussion

This study aimed to resolve intraspecific relationships within the widespread bioindicator diatom species, *Nitzschia palea*. There is considerable overlap in the morphological characters of the varieties described from waters with different pollution levels (*N. palea* and *N. palea* var. *debilis*), making the species complex taxonomically challenging (Trobajo and Cox 2006, Trobajo et al. 2009). Using a combination of morphometrics and phylotranscriptomics, we resolved the *N. palea* species complex into three monophyletic groups. None of these clades contained any strains that could be assigned to *N. palea* var. *debilis* based on the criteria used in Trobajo et al. (2009). However, two narrow strains without deformities (DCG0751 and TC13903) and the strain with the largest mean stria density (TCC13901) were recovered at the linear-lanceolate end of the morphospace in shape outline analysis. These strains partly meet the criteria or fall closest to the ranges given by Trobajo et al. (2009). Both species trees recovered two linear-lanceolate strains (DCG0751 and TCC13903) and TCC907 in Clade B with maximum support values, though as discussed in more detail below, TCC907 is likely of recent hybrid origin. The third linear-lanceolate strain (TCC13901) was recovered outside of the three main clades in both species trees and has the largest mean stria density, providing additional evidence for both morphological and phylogenetic differentiation of this strain (Figure 2, Table S2). Strain TCC13903 in Clade B is capable of self-reproduction, indicating that there is variation in the reproductive characteristics within the *N. palea* species complex, as demonstrated for other diatom species as well (Round et al. 1990).

The remaining six *N. palea* strains fell into two distinct clades (Figure 4, Clades A and C). The morphometric measurements for strains in both clades agree well with the ranges indicated for the nominate variety of *N. palea* (Table S2), which includes populations with medium to large valve width and are most abundant in more polluted freshwater habitats (Trobajo and Cox 2006, Hofmann and Werum 2017). In the shape outline analysis, we recovered these strains at the lanceolate end of morphospace, though they overlap slightly with the linear-lanceolate Clade B strains (PC1, Figure 3). Three strains in Clade C represent sympatric clones isolated simultaneously from a wastewater treatment plant in Belgium, and these are members of one mating type recognized previously as ‘*N. palea* mating group 1’ by Trobajo et al. (2009). Our striae and fibulae measurements for these strains are slightly higher than those given by Trobajo et al. (2009), possibly due to the effects of size reduction in culture (Table S2). On the gradient of valve end shapes, three Clade C strains had the most subcapitate ends in our sample set based on their mean shapes (PC2, Figure 3). However, all three Clade A strains had many deformed valves and missing data, so it is not easy to compare these two groups based on morphometric data alone. Moreover, broader morphometric ranges for the characters measured here have been recorded from natural *N. palea* populations (Bagmet et al. 2020), highlighting that the full range of morphological variation present in this species complex was not represented by the limited number of strains in this study, and many more populations need to be sampled for discussion of *N. palea* morphology. Nonetheless, our species tree estimations recovered ‘N. *palea* mating group 1’ as sister to the common ancestor of Clade A and Clade B, while the remaining three lanceolate strains (Clade A) were recovered as sister to the linear-lanceolate Clade B strains, with maximum support values for both relationships (Figure 4). In conclusion, in agreement with the previous studies, the clades that we recovered did not show a clear morphological distinction as one linear-lanceolate strain (TCC13901) was recovered as a distinct lineage from the linear-lanceolate Clade B, and the two lanceolate clades (A and C) were not monophyletic.

When reconstructing the evolutionary history of a lineage, ILS and gene flow are two main biological causes of incongruence between the species tree and underlying gene trees. ILS occurs when lineages have not been reproductively isolated for long enough for ancestral polymorphisms to fully assort in the descendant lineages so that the gene trees are not reciprocally monophyletic and the speciation history cannot be reflected accurately (Avise 1990). For very recently diverged species, such as the populations of *N. palea* studied here, the probability of ILS is higher due to the persistence of ancestral polymorphisms that can reduce phylogenetic signal (Maddison 1997). Unlike concatenation-based species tree estimation, gene tree summary methods are more robust to discordance between gene trees and species trees in the presence of ILS (Mirarab et al. 2014). Although branch support measures obtained by gene tree summary methods provide useful information, they do not discriminate between the different sources of discordance. Therefore, capturing the underlying agreement and disagreement of gene trees in large phylogenomic datasets requires the use of additional methods (Kumar et al. 2012, Minh et al. 2020b). Although we found evidence of widespread ILS in the evolutionary history of the *N. palea* species complex using gene concordance factors, the estimated species trees showed very high support for all branches, except for one internal branch (Figure 4, Clade C). A large number of gene trees provided evidence of ILS for Clade C, which is expected as these are members of a single interfertile population. Our findings highlight that complementary approaches to gene tree summary methods are necessary to quantify the extent of discordance in the evolutionary history of recently diverged species complexes.

Gene flow among lineages due to hybridization and subsequent gene flow is another process that can generate gene tree discordance and distort phylogenetic signal across the genome (Slatkin and Maddison 1989). Phylogenomic analyses of gene flow have to account for ILS as both processes can generate discordance between gene trees and species trees (Hibbins and Hahn 2022). Models of a strictly bifurcating tree can be rejected in these methods in favor of a network that allows reticulation to summarize instances of gene flow (Wen et al. 2018). Using phylogenetic network methods, we obtained further insights into the relationships between the three clades recovered in the species trees, revealing reticulated evolutionary patterns between lanceolate Clade A and linear-lanceolate Clade B populations (Figure 5B). Although we recovered a clear phylogenetic structure with high statistical support, the mating systems (e.g., cell recognition) in these two lineages apparently have not diverged enough to prevent gene flow. Further sampling from natural populations will show the full extent of gene flow in this species complex and whether there is, for example, a biogeographic component. Moreover, we found strong evidence using phylogenetic networks for the presence of a putative hybrid strain (TCC907) (Figures 4 and 5). Hybridization between intraspecific diatom lineages has been reported for *Seminavis robusta* D.B.Danielidis & D.G.Mann (Decker et al. 2018) and *Eunotia bilunaris* (Ehrenberg) Mills (Vanormelingen et al. 2008) based on evidence from crossing experiments, and between two varieties of *Pseudo-nitzschia pungens* (Grunow ex Cleve) Hasle based on evidence from morphological and phylogenetic analyses (Casteleyn et al. 2009). This study, on the other hand, represents the first report of hybrid detection in a diatom species complex based solely on genome-scale data. Although the parentage cannot be conclusively identified for this putative hybrid given our current sampling, its genome is presumed to be allodiploid, as the inheritance probabilities from Clade A and DCG0751 were almost equal. Evidence for allopolyploidy has also been reported in the diatom *Fistulifera solaris* Mayama, Matsumoto, Nemoto & Tanaka based on genome-wide heterozygosity patterns (Tanaka et al. 2015). Our result was further corroborated with the ABBA-BABA test, which captures the signal for gene flow by comparing parsimony-informative sites that support a different phylogeny than the species tree. DNA sequencing has led to the splitting of once broadly defined widespread diatom taxa into several species; however, our findings indicate that gene flow might obscure species boundaries in some of these, if they have only recently diverged (some species complexes are clearly quite old, e.g., the *Sellaphora pupula* complex: Evans et al. 2008, p. 227).

In conclusion, our analyses revealed a complex evolutionary history of the *N. palea* species complex. We resolved the species complex into three clades, one of which corresponded to a group of narrow linear-lanceolate strains. However, one other linear-lanceolate strain was recovered outside of the three main clades in the phylogenetic analyses, indicating that the linear-lanceolate morphology might have originated multiple times in the evolutionary history of the species. ILS was prevalent across the species tree, providing evidence for the recent divergence of the species complex (Bagley et al. 2020, Kandziora et al. 2021). Moreover, we identified a putative hybrid resulting from recent gene flow between lineages with different morphologies. As demonstrated here, analysis of hundreds of unlinked nuclear loci using summary-coalescence methods and phylogenetic networks that are designed to deal with reticulated patterns, coupled with specific tests to discern a null ILS model and gene flow, can reveal patterns of reticulation and provide crucial information on the relationships of taxonomically challenging diatom lineages.

## Acknowledgments

The authors would like to thank Bertie Joan van Heuven for assistance with SEM imaging; Danny Duijsings for help with supervision, project administration, funding acquisition and logistics; Frederic Rimet for helpful information about cultural strains; and Ekin Çiftçi for help with illustrations. This study was funded by the European Union’s Horizon 2020 research and innovation programme under H2020 MSCA-ITN-ETN grant agreement No 765000 Plant.ID.

## Author contributions

**O. Çiftçi**: Conceptualization, Investigation, Methodology, Visualization, Formal analysis, Software, Writing - original draft. **A.J. Alverson**: Investigation, Resources, Writing - review and editing. **P. van Bodegom**: Supervision, Writing - review and editing. **W. R. Roberts**: Formal analysis, Software, Writing - review and editing. **A. Mertens**: Investigation, Methodology. **B. Van de Vijver**: Conceptualization. **R. Trobajo**: Investigation, Methodology, Writing - review and editing. **D. Mann**: Investigation, Methodology, Writing - review and editing. **W. Piruvano**: Supervision, Project administration. **I. van Eijk**: Investigation, Formal analysis. **B. Gravendeel:** Supervision, Project administration, Funding acquisition, Writing - review and editing.

## References

Allman, E. S., Baños, H. & Rhodes, J. A. 2019. NANUQ: a method for inferring species networks from gene trees under the coalescent model. Algorithms Mol. Biol. 14:24.

Alverson, A. J. 2008. Molecular systematics and the diatom species. Protist. 159:339–353.

Avise, J. C. 1990. Principles of genealogical concordance in species concepts and biological taxonomy. Oxford surveys in evolutionary biology. 7:45–67.

Bagley, J. C., Uribe-Convers, S., Carlsen, M. M. & Muchhala, N. 2020. Utility of targeted sequence capture for phylogenomics in rapid, recent angiosperm radiations: Neotropical Burmeistera bellflowers as a case study. Mol. Phylogenet. Evol. 152:106769.

Bagmet, V. B., Abdullin, S. R., Mazina, S. E., Nikulin, A. Y., Nikulin, V. Y. & Gontcharov, A. A. 2020. Life cycle of Nitzschia palea (Kützing) W. Smith (Bacillariophyta). Russ. J. Dev. Biol. 51:106–114.

Beszteri, B., Ács, É. & Medlin, L. K. 2005. Ribosomal DNA sequence variation among sympatric strains of the Cyclotella meneghiniana complex (Bacillariophyceae) reveals cryptic diversity. Protist. 156:317–333.

Blackmon, H. & Adams, R. A. 2015. EvobiR: tools for comparative analyses and teaching evolutionary biology. Zenodo.

Bolger, A. M., Lohse, M. & Usadel, B. 2014. Trimmomatic: a flexible trimmer for Illumina sequence data. Bioinformatics. 30:2114–2120.

Bonhomme, V., Picq, S., Gaucherel, C. & Claude, J. 2014. Momocs: outline analysis using R. J. Stat. Softw. 56:1–24.

Brown, J. W., Walker, J. F. & Smith, S. A. 2017. Phyx: phylogenetic tools for unix. Bioinformatics. 33:1886–1888.

Casteleyn, G., Adams, N. G., Vanormelingen, P., Debeer, A.-E., Sabbe, K. & Vyverman, W. 2009. Natural hybrids in the marine diatom Pseudo-nitzschia pungens (Bacillariophyceae): genetic and morphological evidence. Protist. 160:343–354.

CEN. 2004. Water quality–Guidance Standard for the Identification, Enumeration and Interpretation of Benthic Diatom Samples from Running Waters. EN 14407:2004.

Cheon, S., Zhang, J. & Park, C. 2020. Is phylotranscriptomics as reliable as phylogenomics? Mol. Biol. Evol. 37:3672–3683.

Cox, M. P., Peterson, D. A. & Biggs, P. J. 2010. SolexaQA: At-a-glance quality assessment of Illumina second-generation sequencing data. BMC Bioinformatics. 11:485.

Decker, S. de, Vanormelingen, P., Pinseel, E., Sefbom, J., Audoor, S., Sabbe, K. & Vyverman, W. 2018. Incomplete reproductive isolation between genetically distinct sympatric clades of the pennate model diatom Seminavis robusta. Protist. 169:569–583.

Degnan, J. H. & Rosenberg, N. A. 2006. Discordance of species trees with their most likely gene trees. Plos. Genet. 2:e68.

Durand, E. Y., Patterson, N., Reich, D. & Slatkin, M. 2011. Testing for ancient admixture between closely related populations. Mol. Biol. Evol. 28:2239–2252.

El-Gebali, S., Mistry, J., Bateman, A., Eddy, S. R., Luciani, A., Potter, S. C., Qureshi, M., … & Finn, R. D. 2019. The Pfam protein families database in 2019. Nucleic Acids Res. 47:D427–D432.

Emms, D. M. & Kelly, S. 2015. OrthoFinder: solving fundamental biases in whole genome comparisons dramatically improves orthogroup inference accuracy. Genome Biol. 16:157.

Emms, D. M. & Kelly, S. 2019. OrthoFinder: phylogenetic orthology inference for comparative genomics. Genome Biol. 20:238.

Evans, K. M., Wortley, A. H., Simpson, G. E., Chepurnov, V. A. & Mann, D. G. 2008. A molecular systematic approach to explore diversity within the Sellaphora pupula species complex (Bacillariophyta). J. Phycol. 44:215–231.

Finlay, B. J., Monaghan, E. B. & Maberly, S. C. 2002. Hypothesis: the rate and scale of dispersal of freshwater diatom species is a function of their global abundance. Protist. 153:261–273.

Finn, R. D., Clements, J. & Eddy, S. R. 2011. HMMER web server: interactive sequence similarity searching. Nucleic Acids Res. 39:W29–37.

Fritz, S. C., Juggins, S., Battarbee, R. W. & Engstrom, D. R. 1991. Reconstruction of past changes in salinity and climate using a diatom-based transfer function. Nature. 352:706–708.

Grabherr, M. G., Haas, B. J., Yassour, M., Levin, J. Z., Thompson, D. A., Amit, I., Adiconis, X., … & Regev, A. 2011. Full-length transcriptome assembly from RNA-Seq data without a reference genome. Nat. Biotechnol. 29:644–652.

Guillard, R. & Lorenzen, C. J. 1972. Yellow-green algae with chlorophyllide c. J. Phycol. 8:10–14.

Haas, B. 2020. TransDecoder (find coding regions within transcripts), Version: 5.5.0. Available at: https://anaconda.org/bioconda/transdecoder/.

Hibbins, M. S. & Hahn, M. W. 2022. Phylogenomic approaches to detecting and characterizing introgression. Genetics. 220:iyab173.

Hofmann, G. & Werum, M. 2017. Freshwater benthic diatoms of Central Europe: Over 800 common species used in ecological assessment. English edition with updated taxonomy and added species. Koeltz Botanical Books, Schmitten-Oberreifenberg Germany, 942 pp.

Huson, D. H., Klöpper, T., Lockhart, P. J. & Steel, M. A. 2005. Reconstruction of Reticulate Networks from Gene Trees. In Miyano, S., Kasif, S., Istrail, S., Pevzner, P. A. & Waterman, M. [Eds.] Research in computational molecular biology. Lecture Notes in Computer Science, Vol. 3500. Springer, Berlin, pp. 233–249.

Huson, D. H. & Bryant, D. 2006. Application of phylogenetic networks in evolutionary studies. Mol. Biol. Evol. 23:254–267.

Huson, D. H., Richter, D. C., Rausch, C., Dezulian, T., Franz, M. & Rupp, R. 2007. Dendroscope: An interactive viewer for large phylogenetic trees. BMC Bioinformatics. 8:460.

Kamikawa, R., Azuma, T., Ishii, K.-I., Matsuno, Y. & Miyashita, H. 2018. Diversity of organellar genomes in non-photosynthetic diatoms. Protist. 169:351–361.

Kandziora, M., Sklenář, P., Kolář, F. & Schmickl, R. 2021. How to tackle phylogenetic discordance in recent and rapidly radiating groups? Developing a workflow using Loricaria (Asteraceae) as an example. Front. Plant. Sci. 12:765719.

Katoh, K. & Standley, D. M. 2013. MAFFT multiple sequence alignment software version 7: improvements in performance and usability. Mol. Biol. Evol. 30:772–780.

Kermarrec, L., Bouchez, A., Rimet, F. & Humbert, J. F. 2013. First evidence of the existence of semi-cryptic species and of a phylogeographic structure in the Gomphonema parvulum (Kützing) Kützing complex (Bacillariophyta). Protist. 164:686–705.

Kloster, M., Kauer, G. & Beszteri, B. 2014. SHERPA: an image segmentation and outline feature extraction tool for diatoms and other objects. BMC Bioinformatics. 15:218.

Krasovec, M., Sanchez-Brosseau, S. & Piganeau, G. 2019. First estimation of the spontaneous mutation rate in diatoms. Genome Biol. Evol. 11:1829–1837.

Kumar, S., Filipski, A. J., Battistuzzi, F. U., Kosakovsky Pond, S. L. & Tamura, K. 2012. Statistics and truth in phylogenomics. Mol. Biol. Evol. 29:457–472.

Lange-Bertalot, H. 1980. New species, combinations and synonyms in the genus Nitzschia. Bacillaria. 3:41–77.

Langmead, B. & Salzberg, S. L. 2012. Fast gapped-read alignment with Bowtie 2. Nat. Methods. 9:357–359.

Lewis, R. J., Jensen, S. I., DeNicola, D. M., Miller, V. I., Hoagland, K. D. & Ernst, S. G. 1997. Genetic variation in the diatom Fragilaria capucina (Fragilariaceae) along a latitudinal gradient across North America. Plant Syst. Evol. 204:99–108.

Li, H. 2015. BFC: correcting Illumina sequencing errors. Bioinformatics. 31:2885–2887.

Li, W. & Godzik, A. 2006. Cd-hit: a fast program for clustering and comparing large sets of protein or nucleotide sequences. Bioinformatics. 22:1658–1659.

Maddison, W. P. 1997. Gene trees in species trees. Syst. Biol. 46:523–536.

Mai, U. & Mirarab, S. 2018. TreeShrink: fast and accurate detection of outlier long branches in collections of phylogenetic trees. BMC Genomics. 19:272.

Mallet, J. 2005. Hybridization as an invasion of the genome. Trends Ecol. Evol. 20:229–237.

Mann, D. G. 1999. The species concept in diatoms. Phycologia. 38:437–495.

Mann, D. G., Trobajo, R., Sato, S., Li, C., Witkowski, A., Rimet, F., Ashworth, M. P., … & Theriot, E. C. 2021. Ripe for reassessment: A synthesis of available molecular data for the speciose diatom family Bacillariaceae. Mol. Phylogenet. Evol. 158:106985.

Minh, B. Q., Schmidt, H. A., Chernomor, O., Schrempf, D., Woodhams, M. D., Von Haeseler, A. & Lanfear, R. 2020a. IQ-TREE 2: New models and efficient methods for phylogenetic inference in the genomic era. Mol. Biol. Evol. 37:1530–1534.

Minh, B. Q., Hahn, M. W. & Lanfear, R. 2020b. New methods to calculate concordance factors for phylogenomic datasets. Mol. Biol. Evol. 37:2727–2733.

Mirarab, S., Reaz, R., Bayzid, M. S., Zimmermann, T., Swenson, M. S. & Warnow, T. 2014. ASTRAL: genome-scale coalescent-based species tree estimation. Bioinformatics. 30:i541–i548.

Pappas, J. L., Kociolek, J. P. & Stoermer E.F. 2014. Quantitative morphometric methods in diatom research. Nova Hedwigia. 143:281–306.

Paradis, E., Claude, J. & Strimmer, K. 2004. APE: analyses of phylogenetics and evolution in R language. Bioinformatics. 20:289–290.

Parks, M. B., Nakov, T., Ruck, E. C., Wickett, N. J. & Alverson, A. J. 2018. Phylogenomics reveals an extensive history of genome duplication in diatoms (Bacillariophyta). Am. J. Bot. 105:330–347.

Patro, R., Duggal, G., Love, M. I., Irizarry, R. A. & Kingsford, C. 2017. Salmon provides fast and bias-aware quantification of transcript expression using dual-phase inference. Nat. Methods. 14:417–419.

Pinseel, E., Kulichová, J., Scharfen, V., Urbánková, P., Van de Vijver, B. & Vyverman, W. 2019. Extensive cryptic diversity in the terrestrial diatom Pinnularia borealis (Bacillariophyceae). Protist. 170:121–140.

Potapova, M. & Hamilton, P. B. 2007. Morphological and ecological variation within the Achnanthidium minutissimum (Bacillariophyceae) species complex. J. Phycol. 43:561–575.

Quijano-Scheggia, S. I., Garcés, E., Lundholm, N., Moestrup, Ø., Andree, K. & Camp, J. 2009. Morphology, physiology, molecular phylogeny and sexual compatibility of the cryptic Pseudo-nitzschia delicatissima complex (Bacillariophyta), including the description of P. arenysensis sp. nov. Phycologia. 48:492–509.

Rhodes, J. A., Baños, H., Mitchell, J. D. & Allman, E. S. 2021. MSCquartets 1.0: quartet methods for species trees and networks under the multispecies coalescent model in R. Bioinformatics. 37:1766–1768.

Rimet, F., Trobajo, R., Mann, D. G., Kermarrec, L., Franc, A., Domaizon, I. & Bouchez, A. 2014. When is sampling complete? The effects of geographical range and marker choice on perceived diversity in Nitzschia palea (Bacillariophyta). Protist. 165:245–259.

Round, F. E., Crawford, R. M. & Mann, D. G. 1990. The Diatoms: Biology and Morphology of the Genera. Cambridge University Press, Cambridge, 747 pp.

Sabater, S. 2000. Diatom communities as indicators of environmental stress in the Guadiamar River, SW. Spain, following a major mine tailings spill. J. Appl. Phycol. 12:113–124.

Sarno, D., Kooistra, W. H. C. F., Medlin, L. K., Percopo, I. & Zingone, A. 2005. Diversity in the genus Skeletonema (Bacillariophyceae). II. An assessment of the taxonomy of S. costatum-like species with the description of four new species. J. Phycol. 41:151–176.

Schindelin, J., Arganda-Carreras, I., Frise, E., Kaynig, V., Longair, M., Pietzsch, T., Preibisch, S., … & Cardona, A. 2012. Fiji: an open-source platform for biological-image analysis. Nat. Methods. 9:676–682.

Slatkin, M. & Maddison, W. P. 1989. A cladistic measure of gene flow inferred from the phylogenies of alleles. Genetics. 123:603–613.

Smol, J. P. & Stoermer, E. F. 2010. The Diatoms: Applications for the Environmental and Earth Sciences. 2nd ed. Cambridge University Press, New York, 667 pp.

Stamatakis, A. 2014. RAxML version 8: a tool for phylogenetic analysis and post-analysis of large phylogenies. Bioinformatics. 30:1312–1313.

Sunagawa, S., Coelho, L. P., Chaffron, S., Kultima, J. R., Labadie, K., Salazar, G., Djahanschiri, B., … & Bork, P. 2015. Ocean plankton. Structure and function of the global ocean microbiome. Science. 348:1261359.

Suyama, M., Torrents, D. & Bork, P. 2006. PAL2NAL: robust conversion of protein sequence alignments into the corresponding codon alignments. Nucleic Acids Res. 34:W609–W612.

Tanaka, T., Maeda, Y., Veluchamy, A., Tanaka, M., Abida, H., Maréchal, E., Bowler, C., … & Fujibuchi, W. 2015. Oil accumulation by the oleaginous diatom Fistulifera solaris as revealed by the genome and transcriptome. Plant Cell. 27:162–176.

The Uniprot Consortium 2019. UniProt: a worldwide hub of protein knowledge. Nucleic Acids Res. 47:D506–D515.

Theriot, E. & Ladewski, T. B. 1986. Morphometric analysis of shape of specimens from the neotype of Tabellaria flocculosa (Bacillariophyceae). Am. J. Bot. 73:224–229.

Theriot, E. C. 1992. Clusters, species concepts, and morphological evolution of diatoms. Syst. Biol. 41:141–157.

Trobajo, R. & Cox, E. J. 2006. Examination of the type material of Nitzschia frustulum, N. palea and N. palea var. debilis. 18th International Diatom Symposium. Biopress Limited.

Trobajo, R., Clavero, E., Chepurnov, V. A., Sabbe, K., Mann, D. G., Ishihara, S. & Cox, E. J. 2009. Morphological, genetic and mating diversity within the widespread bioindicator Nitzschia palea (Bacillariophyceae). Phycologia. 48:443–459.

Trobajo, R., Mann, D. G., Clavero, E., Evans, K. M., Vanormelingen, P. & McGregor, R. C. 2010. The use of partial cox 1, rbc L and LSU rDNA sequences for phylogenetics and species identification within the Nitzschia palea species complex (Bacillariophyceae). Eur. J. Phycol. 45:413–425.

Van Dam, H., Mertens, A. & Sinkeldam, J. 1994. A coded checklist and ecological indicator values of freshwater diatoms from The Netherlands. Netherlands Journal of Aquatic Ecology. 28:117–133.

Vanormelingen, P., Chepurnov, V. A., Mann, D. G., Sabbe, K. & Vyverman, W. 2008. Genetic divergence and reproductive barriers among morphologically heterogeneous sympatric clones of Eunotia bilunaris sensu lato (Bacillariophyta). Protist. 159:73–90.

Waterhouse, R. M., Seppey, M., Simão, F. A., Manni, M., Ioannidis, P., Klioutchnikov, G., Kriventseva, E. V., … & Zdobnov, E. M. 2018. BUSCO applications from quality assessments to gene prediction and phylogenomics. Mol. Biol. Evol. 35:543–548.

Wen, D., Yu, Y., Zhu, J. & Nakhleh, L. 2018. Inferring phylogenetic networks using PhyloNet. Syst. Biol. 67:735–740.

Wickham, H. & Chang, W. 2016. ggplot2: Create elegant data visualisations using the grammar of graphics, Version: 3.3.5. Springer, Switzerland.

Wishkerman, A. & Hamilton, P. B. 2018. Shape outline extraction software (DiaOutline) for elliptic Fourier analysis application in morphometric studies. Appl. Plant. Sci. 6:e01204.

Yang, Y. & Smith, S. A. 2014. Orthology inference in nonmodel organisms using transcriptomes and low-coverage genomes: improving accuracy and matrix occupancy for phylogenomics. Mol. Biol. Evol. 31:3081–3092.

Yu, G., Smith, D. K., Zhu, H., Guan, Y. & Lam, T. T.-Y. 2017. ggtree: an r package for visualization and annotation of phylogenetic trees with their covariates and other associated data. Methods Ecol. Evol. 8:28–36.

Yu, Y. & Nakhleh, L. 2015. A maximum pseudo-likelihood approach for phylogenetic networks. BMC Genomics. 16:S10.

Zhang, C., Sayyari, E. & Mirarab, S. 2017. ASTRAL-III: increased scalability and impacts of contracting low support branches. In RECOMB international workshop on comparative genomics. Springer, Cham, Switzerland, pp. 53–75.

