## Supplemental Figure 1 for "Phylotranscriptomics Reveals the Reticulate Evolutionary History of a Widespread Diatom Species Complex"

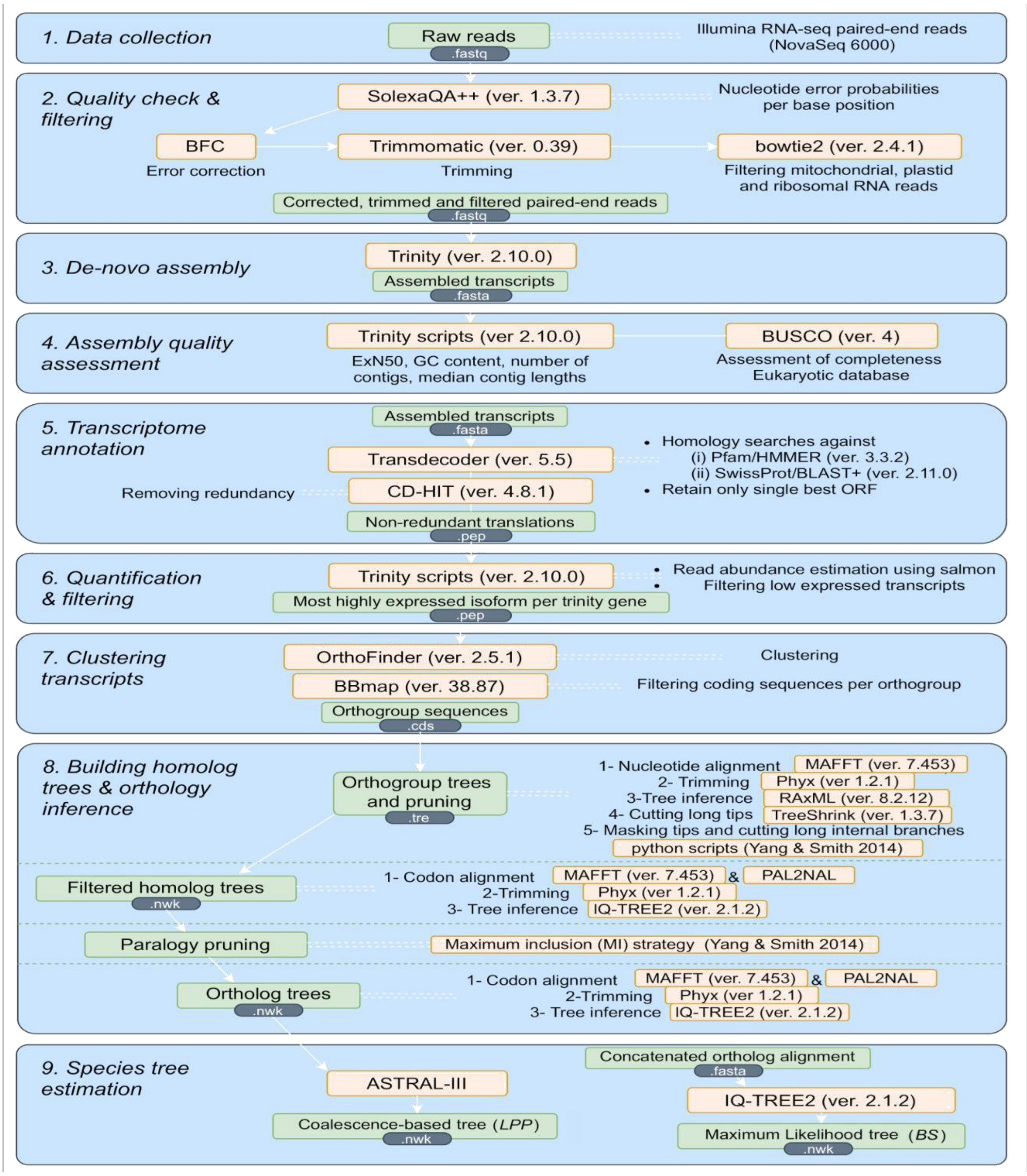


Figure S1. Summary of bioinformatic workflow. Green boxes show the input and output of each step with the file format specified below. Light orange boxes follow the tools and information on the version used.
