## Supplemental Table 1 for "Phylotranscriptomics Reveals the Reticulate Evolutionary History of a Widespread Diatom Species Complex"

Table S1. The number of valves used in shape outline analysis of *Nitzschia palea* per strain and the number of valves removed from the dataset due to tilting caused by stacking on SEM stubs and valve outline deformations.

| Strain identifier | Origin | # of outlines analyzed | # of tilted valves | # of deformed outlines |
| --- | --- | --- | --- | --- |
| DCG0091 | Belgium | 43 | 1 | 0 |
| DCG0092 | Belgium | 50 | 0 | 0 |
| DCG0094 | Belgium | 46 | 1 | 0 |
| DCG0751 | Belgium | 48 | 1 | 0 |
| TCC13901 | France | 50 | 2 | 0 |
| TCC13903 | France | 52 | 1 | 0 |
| TCC523 | Réunion | 51 | 0 | 0 |
| TCC641 | Luxembourg | 36 | 2 | 0 |
| TCC852 | Portugal | 38 | 1 | 7 |
| TCC907 | UK | 0 | 5 | 54 |
