## Supplemental Table 2 for "Phylotranscriptomics Reveals the Reticulate Evolutionary History of a Widespread Diatom Species Complex"

Table S2. Mean value and SD of morphometric measurements per strain compared with type slides of *Nitzschia palea*. Strains are ordered based on their width.

| Strain identifier | Origin | Length (µm) | Width (µm) | L - W ratio | Striae  (10µm) | Fibulae (10µm) |
| --- | --- | --- | --- | --- | --- | --- |
| DCG0091 | Belgium | 15.95±1.13 | 3.89±0.27 | 4.12±0.44 | 39.56±1.09 | 16.06±1.86 |
| DCG0094 | Belgium | 12.12±0.87 | 3.68±0.25 | 3.31±0.35 | 40.38±1.29 | 15.82±1.34 |
| DCG0092 | Belgium | 11.92±0.66 | 3.6±0.16 | 3.31±0.24 | 39.2±1.12 | 15.42±1.85 |
| TCC907 | UK | 20.23±2.6 | 3.58±0.27 | 5.69±0.9 | 35.4±0.91 | 16.89±1.54 |
| TCC641 | Luxembourg | 13.15±0.77 | 3.56±0.39 | 3.72±0.36 | 33.69±1.37 | 14.23±1.31 |
| TCC13901 | France | 19.57±1.72 | 3.53±0.35 | 5.61±0.81 | 41.19±0.97 | 16.6±1.64 |
| DCG0751 | Belgium | 25.62±0.69 | 3.49±0.37 | 7.43±0.82 | 38.76±0.92 | 14.08±1.56 |
| TCC523 | Réunion | 10.33±0.41 | 3.33±0.21 | 3.12±0.25 | 40±0.82 | 16.14±1.13 |
| TCC13903 | France | 33.06±11.5 | 3.27±0.35 | 10.33±3.99 | 40.68±0.67 | 14.59±1.4 |
| TCC852 | Portugal | 12.8±1.17 | 3.15±0.35 | 4.12±0.72 | 40.26±1.11 | 16.78±1.53 |
| *N.palea* type* | Germany | 28.06±4.0 | 3.83±0.25 | unknown | 40.55±2.42 | 16.08±1.65 |
| *N. palea* var. *debilis* type* | Germany | 28.5±1.3 | 3.4±0.2 | unknown | 41.4±1.5 | 14.8±1.1 |

*** Trobajo & Cox (2006)
